# Insights into Plant Interactions and the Biogeochemical Role of the Globally Widespread Acidobacteriota Phylum

**DOI:** 10.1101/2023.10.09.561551

**Authors:** Osiel S. Gonçalves, Alexia S. Fernandes, Sumaya M. Tupy, Tauanne G. Ferreira, Luciano N. Almeida, Christopher J. Creevey, Mateus F. Santana

## Abstract

The prevalence and abundance of Acidobacteriota raise concerns about their ecological function and metabolic activity in the environment. Studies have reported the potential of some members of Acidobacteriota to interact with plants and play a significant role in biogeochemical cycles. However, their role in this context has not been extensively studied. Here, we performed a comprehensive genomic analysis of 758 metagenome-assembled genomes (MAGs) and 121 RefSeq genomes of Acidobacteriota. Our analysis revealed a high frequency of plant growth-promoting traits (PGPTs) genes in the *Acidobacteriaceae, Bryobacteraceae, Koribacteraceae*, and *Pyrinomonadaceae* families. These PGPTs include genes involved in nitrogen fixation, phosphorus solubilization, exopolysaccharide production, siderophore production, and plant growth hormone production. Expression of such genes was found to be transcriptionally active in different environments. In addition, we identified numerous carbohydrate-active enzymes and peptidases involved in plant polymer degradation. By applying an in-depth insight into the diversity of the phylum, we expanded previous role in carbon, nitrogen, sulfur, and trace metal cycling. This study underscores the distinct potential ecological roles of each of these taxonomic groups, providing valuable insights for future research.

## 1. Introduction

Acidobacteriota is a highly prevalent bacterial phylum found in various environments such as soils, freshwater, and marine ecosystems worldwide. It was initially discovered in 1991 with the identification of *Acidobacterium capsulatum* from an acidic mineral habitat (Kishimoto et al., 1991). Since then, the phylum’s widespread distribution was revealed by its distinctive 16S rRNA gene sequence (Rheims et al., 1996; Ludwig et al., 1997; Felske et al., 2000). Studies have shown that Acidobacteriota is a phylogenetically diverse group, consisting of 26 major sequence clades or subdivisions (SDs) (Dedysh and Yilmaz, 2018). It is also a geographically widespread and numerically significant component of the soil microbiota (Kuske et al., 1997).

The prevalence and abundance of these microorganisms raise concerns about their ecological function and metabolic activity, particularly in soil environments (Barns et al., 1999; Quaiser et al., 2003; Lee et al., 2008). In soil ecosystems, members of this phylum are suggesting contributing to nutrient cycling and organic matter decomposition (Crits-Christoph et al., 2018; Woodcroft et al., 2018; Chen et al., 2021). They are known for their ability to tolerate a wide range of soil pH levels and have been found to be particularly abundant in acidic soils. Some Acidobacteriota members have been classified as k-strategists, which means they can thrive in settings with limited nutritional availability, slower growth rates, and high tolerance to toxic compounds (Andrews and Harris, 1986; Kielak et al., 2016b).

Acidobacteriota strains from subdivision 1 (SD1) were identified to act as plant growth-promoting bacteria (Kielak et al., 2016b). These strains produced plant growth-promoting traits (PGPTs), which increased the biomass of roots and shoots in *Arabidopsis thaliana* (Kielak et al., 2016b). This was the first report of Acidobacteriota interacting with the plant. Moreover, given their dominance and metabolic activity in the soil, Acidobacteriota is believed to play a significant role in the biogeochemical cycles of rhizosphere soil (Lee et al., 2008). The diversity of Acidobacteriota in various soils and environments indicates their potential significance in plant-microbe interactions. However, their role in this context has not been extensively studied.

Here, we performed a comprehensive genomic analysis of 758 metagenome-assembled genomes (MAGs) from catalog of Earth’s microbiomes (GEM) (Nayfach et al., 2021a) and 121 RefSeq genomes of Acidobacteriota to better understand the plant interactions and the biogeochemical role of this phylum. Using this genomic database, we identified four ecologically distinct taxonomic groups that have the potential to play a direct role in plant growth. These groups possess genes encoding plant growth-promoting traits (PGPTs), such as nitrogen fixation, phosphorus solubilization, exopolysaccharide (EPS) production, siderophore and plant growth hormones production. Furthermore, we uncovered distinct potential ecological roles for each of these taxonomic groups, which can provide valuable insights for future research.

## 2. Material and Methods

### 2.1. Dataset compilation

We retrieved 758 MAGs from the publicly available GEM catalog (https://portal.nersc.gov/GEM/) (Nayfach et al., 2021b) that were taxonomically affiliated with the phylum Acidobacteriota. To gain insights into these MAGs, we employed QUAST v5.0.2102 to calculate fundamental features such as size, GC content, N50 value, and other relevant metrics. In addition to the MAGs, we also acquired 129 genome RefSeq from the National Center for Biotechnology Information (NCBI). These RefSeq were selected for the purpose of comparison and further analysis. By including these additional genomes, we aimed to broaden the scope of our investigation and potentially uncover valuable similarities or differences between the MAGs and the RefSeq.

We employed a two-step approach to obtain biogeographic and environmental metadata. First, we utilized the genome ID information from the GEM catalog and searched a search on the Metagenome Bin Search tool available on Integrated Microbial Genomes and Microbiomes (JGI IMG, https://img.jgi.doe.gov). This allowed us to gather relevant biogeographic and environmental information associated with the MAGs. Additionally, for the RefSeq genomes, we accessed the metadata available on Biosample on NCBI, which provided us with details of the geographical origin and environmental context of these genomes.

### 2.2. Phylogenetic analyses

CheckM v1.0.13 (Parks et al., 2015) with the tree_qa command was employed to extract 43 single-copy, protein-coding marker genes. These marker genes were utilized to assess phylogenetic markers in a dataset comprising 758 assembled metagenomic bins and 121 genomes. The concatenated protein alignments of 43 universal marker genes obtained from CheckM were subsequently used to reconstruct maximum-likelihood phylogenetic tree using IQ-TREE v1.6.11 (Minh et al., 2020). The reconstruction process incorporated specific parameters (‘-m TEST -bb 1000’) to ensure accurate tree generation. The phylogenetic tree was uploaded to iTOL (Letunic and Bork, 2019), where it underwent visual annotation, such as color coding and heatmaps were applied.

### 2.3. Annotations and metabolic predictions

MAG gene prediction was determined using Prokka v1.14.5 (Seemann, 2014) with specific parameters (‘—metagenome’) then annotated by KOfam and custom HMM profiles within METABOLIC v.4.0 (Zhou et al., 2022) and eggNOG-emapper v.2.1.2 (Cantalapiedra et al., 2021) with default settings. Additionally, METABOLIC-G v.4.0 was used to profile metabolic and biogeochemical traits, and functional networks in the Acidobacteriota MAGs and genomes. dbCAN2 (Zhang et al., 2018) within METABOLIC v.4.0 was used to annotate proteins carbohydrate-active enzymes (CAZymes) using default thresholds. Peptidases were searched against MEROPS ‘pepunit’ database (Rawlings et al., 2018) also implemented in METABOLIC. The annotations of genes of interest were compared among the outputs of different annotation tools. Amino acid sequences of selected MAGs were used to detect known prokaryotic antiviral systems using DefenseFinder (https://defense-finder.mdmparis-lab.com) (Tesson et al., 2022).

### 2.4. Identification of plant growth-promoting traits (PGPTs)

We performed alignment of MAGs’ protein sequences to identify genes related to plant growth-promoting traits (PGPTs) within the genomes. This alignment was carried out using a combination of BLASTP and HMMER tools in PGPT-Pred database from PLaBAse (Patz et al., 2021). In addition, to ensure accuracy, the annotations were further validated through BLASTP searches against the NCBI nonredundant protein, RefSeq, and UniprotKB/Swiss-Prot databases. Hits with an E-value <1e−5 were considered as significant for both approaches. To determine the MAGs potential to interact with plant, we established a criterion based on the presence/absence of 86 PGPTs genes involved in nitrogen fixation, phosphorus solubilization, as well as the production of EPSs, siderophore and plant growth hormones. Specifically:

- Nitrogen-fixing genes: *nifA, nifB, nifD, nifE, nifF, nifH, nifHD1, nifHD2, nifJ, nifK, nifM, nifN, nifQ, nifS, nifT, nifU, nifV, nifW, nifX, nifZ*.
- *Exopolysaccharide production: epsE, epsD, epsF, epsH, epsI, epsJ, epsL, epsM, epsN, epsO*.
- Root colonization by nodulation: *nodA, nodB, nodC, node, nodF, nodI, nodJ, nodU, nod, nodT, nolM, noeA, noeB, noeC, noeD, noeE, nodN, nodN_like, nodO, nodP, nodS, nodS_like, nodY, nodZ, nodV, nodV_like, nodW, nodX, nodX, nodO*.
- Oxidative stress|ROS scavenging: *sodN, sodC, sod3*.
- Iron acquisition: *lipA, lipB, lipL, lipL2, lipM, lplA*.
- Salinity stress-potassium transport: *kdpA, kdpB, kdpC, kdpD, kdpE, kdpF*.
- Plant embryogenesis-spermidine: *puuA, puuB, puuC, puuD, puuE, pup*
- IAA related tryptophan metabolism: *trpA, trpB, trpC, trpCF, trpD, trpE, trpEG, trpG, trpDG, trpF, trpS, trpR*

See Supplementary Table 6 for more details. The frequency of PGPT genes was transformed into ln(x) values, and a heatmap was generated using Clustvis v.1.0 (https://biit.cs.ut.ee/clustvis/) (Metsalu and Vilo, 2015). To *nif*-genes analysis of 17 MAGs with role in N2 fixation were selected. Nif-cluster were extracted and syntenic analysis was performed using clinker and clustermap.js (Gilchrist and Chooi, 2021).

### 2.5. Metatranscriptomic analysis

Publicly available metatranscriptomic data from JGI IMG was used to investigate the expression of PGPT genes, as well as metabolic and biogeochemical traits. We specifically searched for Acidobacteriota RNASeq Expression Studies, and *Acidobacteriaceae* bacterium URHE0068 and *Chloracidobacterium thermophilum* B were considered. We selected the studies “*Avena fatua* rhizosphere and bulk microbial communities” (submission ID 49153, 48966, 48929, 49411, 49409, and 49669) and “Grassland soil microbial communities” (submission ID 49669, 49668, 49671, 49675, 49677, 49686) for *Acidobacteriaceae* bacterium URHE0068. In addition, we chose the study “Anoxygenic and chlorotrophic microbial mat microbial communities from Yellowstone National Park” (submission ID 64582, 64581, and 64584) for *C. thermophilum*.

Subsequently, we mapped PGPT genes, metabolic traits, and biogeochemical traits in these selected studies. To quantify gene expression, read counts per transcript were normalized by the total number of samplings reads and the length of each transcript. FPKM values were then calculated using the formula FPKM = (Number of mapped reads for the gene / Length of the gene in KB) / (Total number of mapped reads in the sample / 1 million). This normalization and calculation procedure allowed for the comparison of gene expression levels across different samples and genes of interest.

### 2.6. Data availability

The genomes used in this study are publicly available https://portal.nersc.gov/GEM/. Source data with genes associated with plant growth-promoting traits (PGPTs) in Acidobacteriota MAGs are available in the Zenodo repository: https://doi.org/10.5281/zenodo.7957159. The datasets generated and analyzed during the study are provided with this paper in supplementary information.

## 3. Results

### 3.1. A comprehensive analysis of 758 MAGs reveals the potential for plant growth promotion of Acidobacteriota in diverse environments

To investigate the potential interaction between the Acidobacteriota phylum and plants, we collected a total of 758 MAGs from the GEM catalog (Nayfach et al., 2021a) and 121 RefSeq genomes from NCBI (Supplementary Table 1, Supplementary Table 2), which provided us with a comprehensive dataset to explore the diversity and abundance of Acidobacteriota within different environments. The dataset included MAGs from all continents and oceans on Earth, with a notable emphasis on samples from North America and Europe (Fig. 1A). These MAGs were obtained from diverse environments, including soils and other terrestrial habitats (243), ocean and other aquatic environments (173), plant host-associated (52), and engineered environments (52) (Fig. 1B, Supplementary Table 3). In addition, the sizes of Acidobacteriota MAGs ranged from 1 Mb to 11 Mb, with the largest one discovered in the UBA5066 and Bryobacterales order (Supplementary Table 4, Supplementary Fig 1).

**Figure 1.**
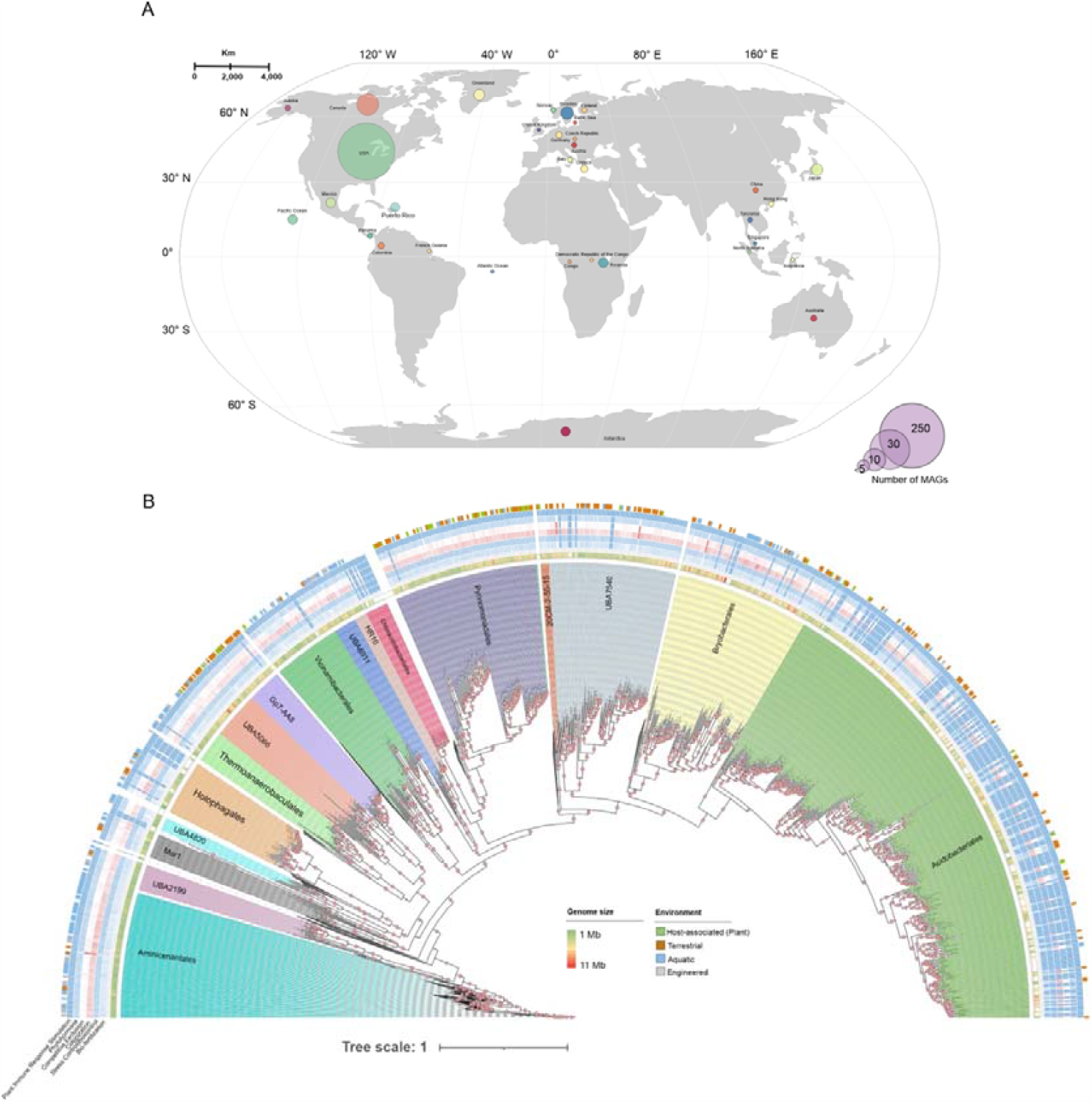
Phylogeny and an overview of their PGPTs potential and global distribution of 885 genomes of Acidobacteriota phylum. (**A**) A maximum likelihood phylogenetic tree of 879 genomes including the 758 metagenome assembled genomes (MAGs) from GEM catalog and 121 RefSeq genomes. The phylogeny is based on 43 single-copy, protein-coding marker genes identified using CheckM. Acidobacteriota families are marked in different background colors. The first arc of the tree displays the environmental source of each genome, followed by sequential arcs representing the relative abundance of genes associated with plant growth-promoting traits (PGPTs), and the last arc represents the genome size of each MAGs. Phylogenetic tree was built under the model of rate heterogeneity LG+F+I+64, and a maximum of 1,000 bootstrap replicates. Bootstraps are shown in pink circles. The tree is drawn to scale, with branch lengths in the same units as those of the evolutionary distances used to infer the phylogenetic tree. (**B**) The global distribution of the Acidobacteriota described in this study in a map generated using ‘ggplot2’ package in R. The abundance is highlighted in distinct circle size.

These MAGs represented 13 different taxonomic classes, with *Acidobacteriaceae* (n= 372) being the most abundant class in the dataset (Supplementary Fig. 1). It is noteworthy that most of the isolates cultivated to date from this class are affiliated with formal SD1, which suggests that this subdivision is the most well-studied group within the Acidobacteriota phylum (Kielak et al., 2016a). Furthermore, we observed that the predominant members of the *Acidobacteriaceae* family were bacteria of the order Acidobacteriales (Supplementary Table 1).

We constructed a phylogenetic tree using 43 single-copy marker genes to understand the evolutionary relationships between Acidobacteriota genomes (Fig. 1B). Our dataset included 758 MAGs obtained from various environments, as well as 121 Acidobacteriota reference genomes publicly available. The phylogenetic tree analysis revealed the formation of 17 distinct branches, thereby the presence of high yet uncharacterized Acidobacteriota species-level diversity (Fig. 1B). These branches represented different taxonomic groups within the dataset, with most genomes (n=211) were found to belong to the Acidobacteriales order, indicating their abundance and widespread distribution in various environments.

Subsequently, we conducted a genome mining analysis on Acidobacteriota MAGs to identify genes associated with PGPTs, to explore the potential of this phylum in plant growth promotion. We categorized these genes into six classes associated with PGPTs, namely bio-fertilization, stress-control/biocontrol, colonization, competitive exclusion, phytohormone production, and plant immune response stimulation (Supplementary Table 5). Our analysis revealed that the colonization and competitive exclusion PGPTs classes had a broad presence in the Acidobacteriota phylum (Fig. 1B), and their genes were linked to indirect effects on plant growth, such as the utilization of amino acids, carbohydrates, and lipids derived from plants, as well as motility and chemotaxis. In general, we noticed that the abundance of these categories varied significantly, particularly within certain taxa such as members of the *Acidobacteriaceae* (Supplementary Fig. 2). Notably, the Acidobacteriales and Holophagales order showed levels of PGPTs associated with bio-fertilization and phytohormone production. These classes are known to play a direct role in plant growth and contain genes involved in iron acquisition, nitrogen assimilation and regulation, mineral solubilization (K, P), auxin and cytokinin synthesis, vitamin production, plant germination, H_2_S production, and metabolism (Supplemental data 1).

After observing that numerous genes within these categories may not be directly contribute to plant growth, we specifically chose 86 PGPTs genes that are involved in nitrogen fixation, phosphorus solubilization, as well as the production of EPSs, siderophore and plant growth hormones (Supplementary Table 6). A clear division into two clusters based on family and gene frequency was observed (Fig. 2). The first cluster comprised the families *Acidobacteriaceae, Bryobacteraceae, Koribacteraceae*, and *Pyrinomonadaceae*, which exhibited a high frequency of PGPT genes. Conversely, *Aminicenantaceae, Chloracidobacteriaceae, Holophagaceae, Thermoanaerobaculacea*, and *Vicinamibacteraceae* displayed lower frequencies of these genes (Supplementary Fig. 3). The genes associated with tryptophan metabolism (*trpABCDEGFS*), potassium solubilization (*kdpABCDEF*), and EPS production (*epsDFHILMNO*) were predominantly identified in the second cluster. Furthermore, we observed a high occurrence of nitrogen-fixing genes in *Acidobacteriaceae* and *Holophagaceae* (Supplementary Table 6, Fig. 2). These findings suggest that the second cluster is potentially involved in a range of functions related to nutrient cycling and soil health.

**Figure 2.**
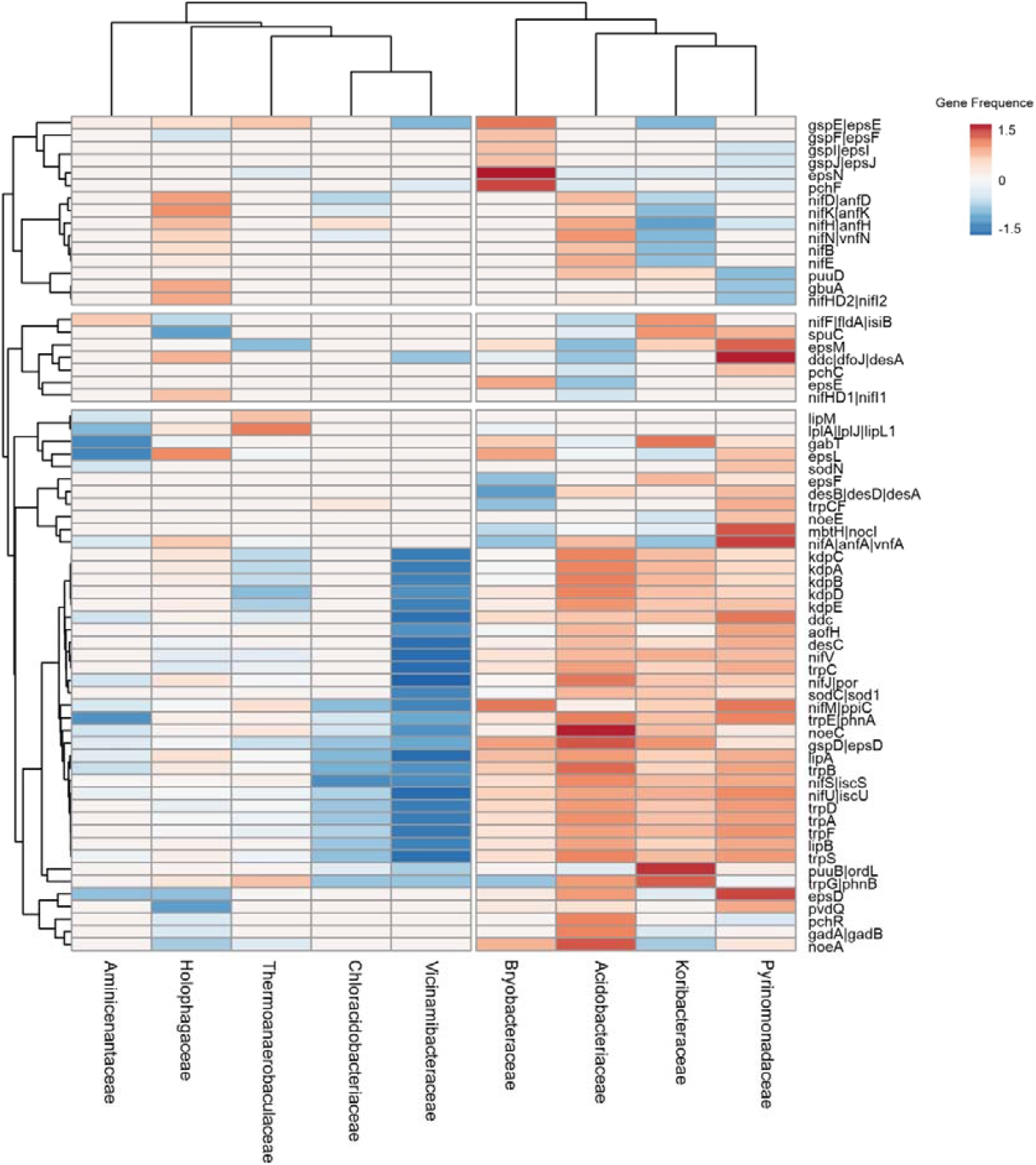
Heatmap depicting the distribution of 86 genes associated with plant growth-promoting traits (PGPTs) that directly contribute to plant growth. The abundance values have been logarithmically transformed (ln(x)-transformed). Row centering and unit variance scaling have been applied to enhance visualization. Missing values were estimated using imputation techniques. The rows and columns have been clustered using correlation distance and average linkage methods.

**Figure 3.**
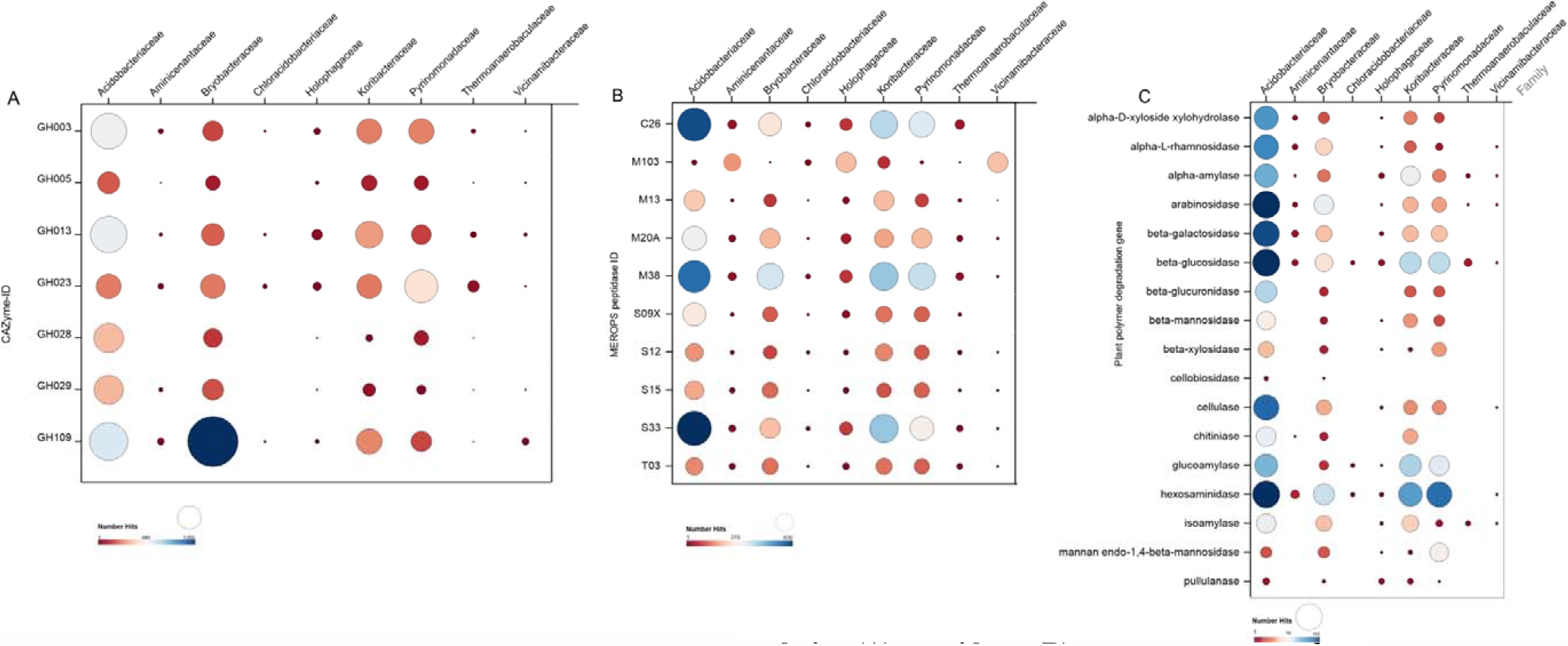
The abundances of genes involved in carbohydrate degradation (A), peptidases (B), and plant polymer degradation (C) across the MAGs. Each circle’s size represents the number of hits for the corresponding gene. The abbreviation “GH” refers to glycoside hydrolases. The x-axis represents the genes, while the y-axis displays the taxonomic family of Acidobacteriota. For a comprehensive description of the carbohydrate-active enzymes (CAZymes) and peptidases present in the genomes, please refer to Supplementary Table 7 and Table 8, respectively.

### 3.2. The potential of Acidobacteriota to act in plant polymer degradation

Carbohydrate Active Enzymes (CAZymes) and peptidases play a crucial role in plant growth and development. These enzymes are responsible for the synthesis, modification, and breakdown of carbohydrates in plants (Drula et al., 2022). Our analysis mapped of 39,722 CAZymes distributed across 758 MAGs of Acidobacteriota (Supplementary Table 7). Among these enzymes, a predominance of GH (Glycosyl Hydrolases) and the presence of some PL (Polysaccharide Lyases) in small number were identified. In particular, the GH enzyme family stood out, representing the seven most abundant CAZymes in these genomes. These enzymes were identified as GH3, GH5, GH13, GH23, GH28, GH29, and GH109, with varying in numbers, ranging from 832 to 4299 Hits, and abundance within the phylum, being *Acidobacteriaceae* and *Bryobacteriaceae* exhibited a higher abundance of these enzymes in their genomes (Supplementary Table 7, Supplementary Fig. 4A, Fig.3A). Particularly noteworthy was the significant presence of GH109 in both families, suggesting their involvement in activities related to α-N-acetylgalactosaminidase. Additionally, *Acidobacteriaceae* showed high quantities of GH3 and GH13 enzymes, indicating their capacity for cellulose, glucan, starch, and peptidoglycan degradation. In addition to these activities, the seven most abundant enzymes also individually contribute to the degradation of mannan, chitooligosaccharides, pectin, and fucose.

A total of 32,566 proteolytic enzymes were mapped, grouped according to their catalytic type, including five families of aspartic peptidases (A), 18 families of cysteine peptidases (C), 62 families of metallo peptidases (M), 2 families of asparagine peptide lyases (N), 1 family of mixed peptidase (P), 37 families of serine peptidases (S), 6 families of threonine peptidases (T), and 3 families of peptidases with unknown catalytic type. Additionally, 761 inhibitors belonging to 5 distinct families were mapped in these MAGs (Supplementary Table 8).

Among the peptidases, the eight most abundant families were selected from the mapped genomes. These families include: C26, corresponding to the family of gamma-glutamyl hydrolases; M103, containing the peptidase TldD; M13, containing metalloendopeptidases with restricted activity to substrates smaller than proteins; M20A, containing carboxypeptidases; M38, corresponding to the family of beta-aspartyl dipeptidases; S09X, containing a diverse set of serine-dependent peptidases; S12, containing serine-type D-Ala-D-Ala carboxypeptidases; S15, containing Xaa-Pro dipeptidyl peptidase; S33, containing exopeptidases that act on the N-terminus of peptides; and T03, which exhibits activities of aminopeptidase and aminotransferase (Fig. 3B). The *Acidobacteriaceae* family hosted most proteolytic enzymes, followed, in smaller quantities, by the *Koribacteriaceae, Pyrinomonadaceae*, and *Bryobacteriaceae* families, respectively (Supplementary Fig. 4B, Fig. 3B).

*Acidobacteriaceae* showed a higher number of enzymes related to plant polymer degradation, such as GH2, GH42, GH13, GH144, GH149, GH6, GH9, GH55, GH77, GH97, GH1, GH23, GH15, GH31, GH10, GH115, GH30, GH67, GH8, GH74, GH120, and GH39 (Supplementary Table 7), indicating the possible involvement of *Acidobacteriaceae* in the degradation of cellulose, pectin, cellobiose, galactose, mannose, glucuronic acid, glucan, and arabinofuranose cleavage (Fig. 3C). Furthermore, the families *Bryobacteraceae, Koribacteraceae*, and *Pyrinomonadaceae* also exhibited a significant quantity of these enzymes. Altogether, this highlights the significant potential of this bacterial family to act in plant polymer degradation.

### 3.3. Role of Acidobacteriota in Biogeochemical Cycles

Our findings highlight the significant role of organisms from the phylum Acidobacteriota in four biogeochemical cycles: carbon, nitrogen, sulfur, and elemental metal cycles (Supplementary Table S9, Fig. 4). We identified 618 MAGs from this phylum that contribute to these reactions, and most of them carry *acs* genes, which are important for acetate oxidation (Fig. 4A). Moreover, we observed a similar pattern of classes participating in the sulfur cycle pathways, which exhibited substantial Acidobacteriota involvement (Supplementary Fig. 5). Within this cycle, we identified eight classes of the Acidobacteriota, comprising a total of 436 MAGs engaged in sulfur oxidation reactions (Fig. 4B). Additionally, in pathways involving the reduction of sulfur, 278 MAGs were found, which shared the same classes present in the oxidation pathways, differing only by the inclusion of subgroup 26. Notably, the MAGs primarily encoded genes associated with sulfide oxidation (*fccB, sqr*), sulfite reduction (*dsrABD, asrABC*), sulfur oxidation (*sdo, sor*), sulfur reduction (*sreABC, sor*), thiosulfate oxidation (*soxBCY*), sulfate reduction (*aprA, sat*), and thiosulfate disproportionation (*aprA*).

**Figure 4.**
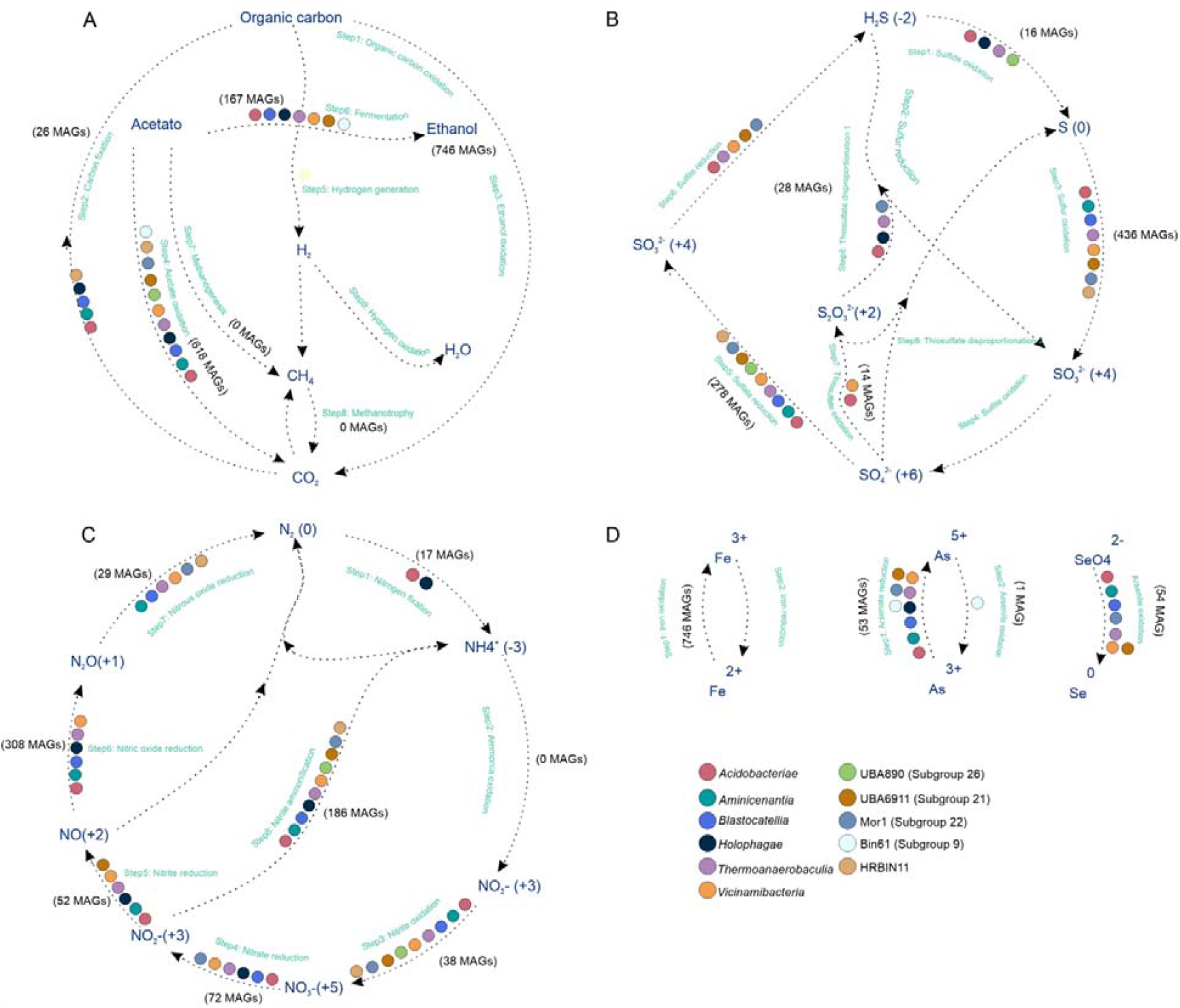
The genomic-based predictions regarding the potential biogeochemical role of Acidobacteriota in various elemental cycles. The taxonomic classes are color-coded to represent their involvement in different steps within the Carbon (**A**), Nitrogen (**B**), Sulfur (**C**), and trace metal (**D**) cycles. The number of Metagenome-Assembled Genomes (MAGs) participating in each element transformation is indicated within parentheses. It is important to note that only pathways encoded in at least one MAG are displayed, and when all MAGs are involved in a particular pathway, taxonomic classes are not color-coded. Each arrow in the figure signifies a single transformation or step within a specific cycle.

Furthermore, a significant contribution of Acidobacteriota to nitrogen balance in the environment (Supplementary Fig. 5). While individuals from all eleven taxonomic classes of the phylum Acidobacteriota participated in eight stages of the nitrogen cycle, they were notably absent in the ammonia oxidation pathway (Fig. 4C). Conversely, in pathways crucial for nitrogen absorption and assimilation by plants and microbiota, such as nitrification (*norBC*) and nitrite ammonification (*nrfADH, nirBD*), a higher number of taxa were predicted to be involved. Specifically, 308 MAGs were associated with nitrification, and 186 MAGs were associated with nitrite ammonification (Fig. 4C). Moreover, we identified 17 MAGs belonging to the classes *Acidobacteriae* and *Holophagae* that possess the capability to engage in biological nitrogen fixation. Marker genes involved in biological nitrogen fixation, namely *nifDK* and *nifH*, were successfully identified in these MAGs.

Lastly, with regard to elemental metals, all 746 MAGs were found to participate in redox reactions involving iron (Fig. 4D). However, the elements selenium and arsenic displayed minimal participation from individuals within the taxonomic classes. Specifically, only one MAG was identified in the arsenite oxidation pathways (Fig. 4D). In summary, our research underscores the crucial role played by Acidobacteriota in important biogeochemical processes. Their involvement spans the carbon, nitrogen, sulfur, and elemental metal cycles, demonstrating their significance in maintaining environmental balance and ecosystem functioning.

### 3.4. Expansion of Nitrogen-fixing activity in Acidobacteriota

Although nitrogen-fixing genes have been acknowledged for their significance (Linta and Xinning, 2022), their distribution and role within the phylum remain largely unexplored. Consequently, additional research is necessary to uncover the prevalence of these genes in various bacterial families and their functional implications in different ecological settings. A search was conducted to identify evolutionary conserved genes associated with nitrogen fixation in the genomes under study. A total of seventeen MAGs were discovered to contain genes linked to nitrogen fixation. Upon analysis, it was determined that the genetic capacity for N_2_ fixation was linked to the following families: *Holophagaceae* (6), *Acidobacteriaceae* (5), *Bryobacteraceae* (5), and *Koribacteraceae* (1). Notably, *Holophagaceae* displayed the highest prevalence, while *Koribacteraceae* exhibited the lowest frequency. At genus level, we identified *Holophaga* (4), *Geothrix* (2), *Terracidiphilus* (1), and *Granulicella* (1). Furthermore, it was observed that 8 MAGs did not belong to any classified genera within the phylum (Supplementary Table S9). The majority of genes were associated with MAGs predominantly distributed in terrestrial environments (15), with a smaller presence in aquatic environments (2) and only one MAG identified in engineered environments (1). Within the terrestrial environment, some MAGs were found in peat soils (2), while the majority were detected in permafrost (10) (Supplementary Fig. 6A).

We mapped eight nif genes directly involved in the N2 fixation, including *nifA* (5), *nifB* (14), *nifD* (17), *nifH* (6), *nifJ* (16), *nifK* (16), *nifS* (12), and *nifW* (4). The *nifD* gene was found in all analyzed genomes, either in its complete form or as fragments. Notably, OTU-17217, affiliated with the *Terracidiphilus* genus, exhibited a higher abundance of nitrogen assimilation genes, as well as other genes associated with the organization of nitrogen fixation operons (Supplementary Fig. 6B) (Garcia et al., 2020). However, contig fragmentation resulted in several reference sequences containing more than one fragment per contig. Overall, our analysis provided a comprehensive understanding of how nitrogen-fixing genes are distributed, their functional implications, and their organization within various ecological settings in the Acidobacteriota phylum.

### 3.5. Evidence of transcriptional activity of PGPTs in metatranscriptomic data

Lastly, we employed metatranscriptomic data from *Acidobacteriaceae* bacterium URHE0068 to investigate their transcriptional activity related to PGPTs (Fig. 5). These data were collected from various environments, including grassland soil microbial communities (Fig. 5A) and *Avena fatua* rhizosphere and bulk microbial communities (Fig. 5B, Supplementary Table S10).

**Figure 5.**
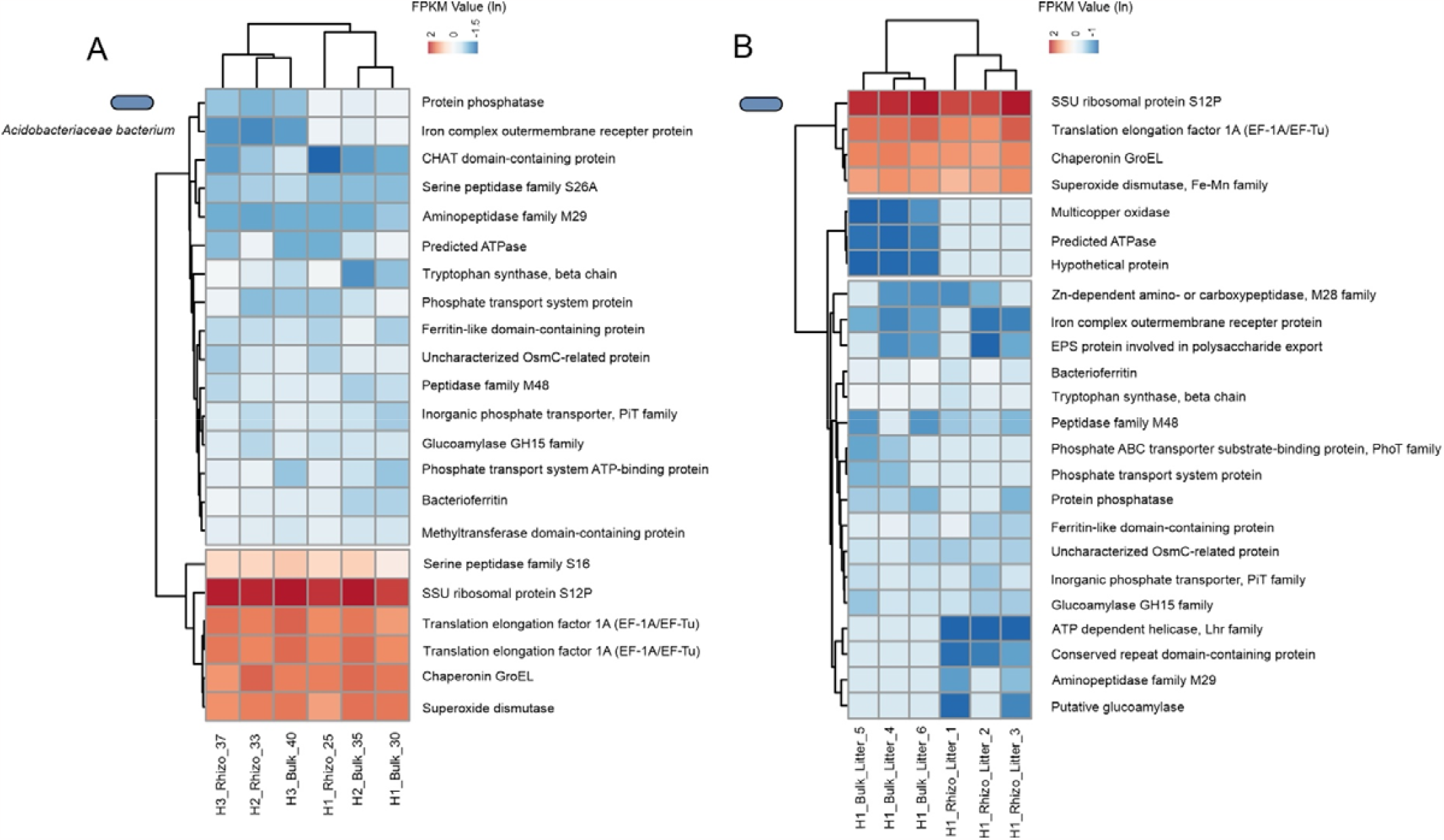
Metatranscriptomic activity of genes associated with PGPTs in two strains of Acidobacteriota across different environmental settings. The expression levels, measured in Fragments Per Kilobase per Million mapped fragments (FPKM), are shown for *Acidobacteriaceae* bacterium URHE0068 in Grassland soil microbial communities (A) and *Avena fatua* rhizosphere and bulk soil microbial communities (B). The expression levels are reported in ln-transformed FPKM. The rows in the figure represent individual samples, without any correlation, and have been clustered using Euclidean distance and average linkage.

Our analysis revealed that both species exhibited metabolic activity in these environments, as evidenced by the high expression levels of general metabolism (Fig. 5). Furthermore, PGPTs associated with phosphorus solubilization, siderophore production, EPS, and genes related to plant growth hormones exhibited above-average relative transcriptional activity, particularly in terrestrial ecosystems associated with plants. Interestingly, several peptidase genes showed the highest levels of transcriptional activity in grassland soil (Fig. 5B, C). Additionally, we observed differential expression of the aminopeptidase family M9 gene between soil associated with root (rhizosphere) and soil not associated (bulk soil) of *A. fatua*. Similarly, phosphorus transport genes exhibited above-average relative transcriptional activity in the rhizosphere of the host plant. Overall, our findings shed light on the transcriptional activity of Acidobacteriota and their involvement in PGPTs. These results highlight their potential contributions to nutrient cycling and plant-microbe interactions in various ecosystems.

## 4. Discussions

Acidobacteriota is a prominent bacterial taxon widely observed in soils worldwide, often constituting a substantial portion of the total bacterial community, reaching up to 52% in some cases (Kuske et al., 1997; Barns et al., 1999; Delgado-Baquerizo et al., 2018). These bacteria are commonly associated with the rhizosphere, the region surrounding plant roots, where intricate interactions between plants and microorganisms occur, mutually influencing each other. Despite their high abundance in this environment, our understanding of how Acidobacteriota members specifically interact with plants and contribute to biogeochemical processes remains limited. Here, we took advantage of metagenome data spanning various environmental contexts within the Acidobacteriota to elucidate the roles played by this phylum in plant-microbe interactions and their impact on the biogeochemical dynamics.

We explored 758 MAGs from GEM catalog and 121 RefSeq genomes of Acidobacteriota from wide range of habitats worldwide, which allowed us to explore the ecological roles and adaptations of this phylum in different environments. Our comprehensive analysis of the metagenome data allowed us to identify a wide distribution of PGPTs genes within the Acidobacteriota phylum. Notably, the colonization and competitive exclusion classes of PGPTs were found to be present in numerous Acidobacteriota members. The colonization PGPT class encompasses genes that are involved in establishing robust interactions between Acidobacteriota and plants. These genes contribute to processes such as adhesion, biofilm formation, and the ability to colonize plant tissues (Compant et al., 2010; Knights et al., 2021; Ishizawa et al., 2022). Similarly, the competitive exclusion PGPT class plays a vital role in plant-microbe interactions. Genes within this class enable Acidobacteriota to outcompete other microorganisms (Ghoul and Mitri, 2016). Previous studies have provided evidence that members of the Acidobacteriota possess multiple conjugative and integrative elements within their genomes and upregulate the expression of specific genes to persistence in the soil environments (Greening et al., 2015; Gonçalves and Santana, 2021).

Our analysis identified four major Acidobacteriota families, namely *Acidobacteriaceae, Bryobacteraceae, Koribacteraceae*, and *Pyrinomonadaceae*, which exhibited a considerable number of 86 keys PGPTs genes. These families have the potential to directly interact with plants, suggesting their ability to promote plant growth and development. The presence of these PGPT genes within these Acidobacteriota families highlights their importance in establishing beneficial relationships with plants and their potential contributions to enhancing plant growth. *Acidobacteriaceae*, a family within the SD1 of the Acidobacteriota phylum, stands out as the most abundant and extensively cultured group within the phylum (Kielak et al., 2016a; Kalam et al., 2020). It is considered the dominant division of Acidobacteriota. The initial discovery of plant growth-promoting abilities within the Acidobacteriota phylum was associated with *Acidobacteriaceae*. Subsequently, another study reported the presence of plant growth-promoting members in two additional families: *Bryobacteraceae* (SD3) and *vicinamibacteraceae* (SD6). Although no typical bacterial plant growth-promoting traits were identified in vitro for these strains, they were found to enhance the growth of duckweed and exhibited the ability to colonize plant surfaces (Yoneda et al., 2021).

We profiled the carbohydrate degradation and peptidase activities of Acidobacteriota. Our analysis revealed that the enzymatic repertoire of Acidobacteriota, being some members encode up to 200 CAZymes and more than 100 peptidases, specifically in the four major families associated with PGPTs, was enriched with genes involved in the degradation of plant polymers. This suggests a potential connection between carbohydrate degradation and PGPTs in Acidobacteriota. However, it is important to consider that this correlation could be influenced by the dataset itself, as the number of MAGs available for these specific Acidobacteriota families may are abundant compared to others.

Acidobacteriota may play a significant role in biogeochemical cycling, although many aspects of their specific contributions are still not well understood (Kalam et al., 2020). Here, we applied an in-depth insight into the diversity of the phylum and to expand previously role in sulphur and nitrogen cycling (Supplementary note 1). Our analysis revealed a significant presence of Acidobacteriota in both the sulfur and nitrogen cycles, which are vital processes in ecosystems, incorporating essential nutrients into the soil through sulfur oxidation and nitrogen fixation (Ågren et al., 2012; Vavourakis et al., 2019), making them available to plants. Finally, our study provided evidence of the metabolic activity of Acidobacteriota members in diverse environments, demonstrating their contribution to the expression of crucial genes involved in PGPTs. These findings shed light on the roles played by Acidobacteriota in plant-microbe interactions and their impact on biogeochemical processes. The transcriptional activity of highly expressed core genes further emphasized the metabolic relevance of Acidobacteriota in different environments.

We aimed to shed light on the role of specific families within Acidobacteriota in plant-microbe interactions and their potential impact on biogeochemical processes that contribute to soil functioning (Fig. 6). We focused on six representatives of Acidobacteriota known to have a more direct relationship with plant interactions. These strains possess the ability to form flagella, enabling them to encounter root exudates and mucilage components (Turnbull et al., 2001). Chemotaxis driven by flagella likely plays a crucial role in root colonization. Additionally, we found a consensus among these members regarding the presence of the high-affinity phosphate transport system *pstSCAB*, located on the outer side of the cell membrane (Fig. 6). This indicates their ability to transport inorganic phosphate (Martin and Liras, 2021). Furthermore, we demonstrated that these bacterial strains may degrade complex plant polymers such as chitin, cellulose, and hemicellulose, as reported in previous studies (Ward et al., 2009; Rawat et al., 2012; Belova et al., 2018).

**Figure 6.**
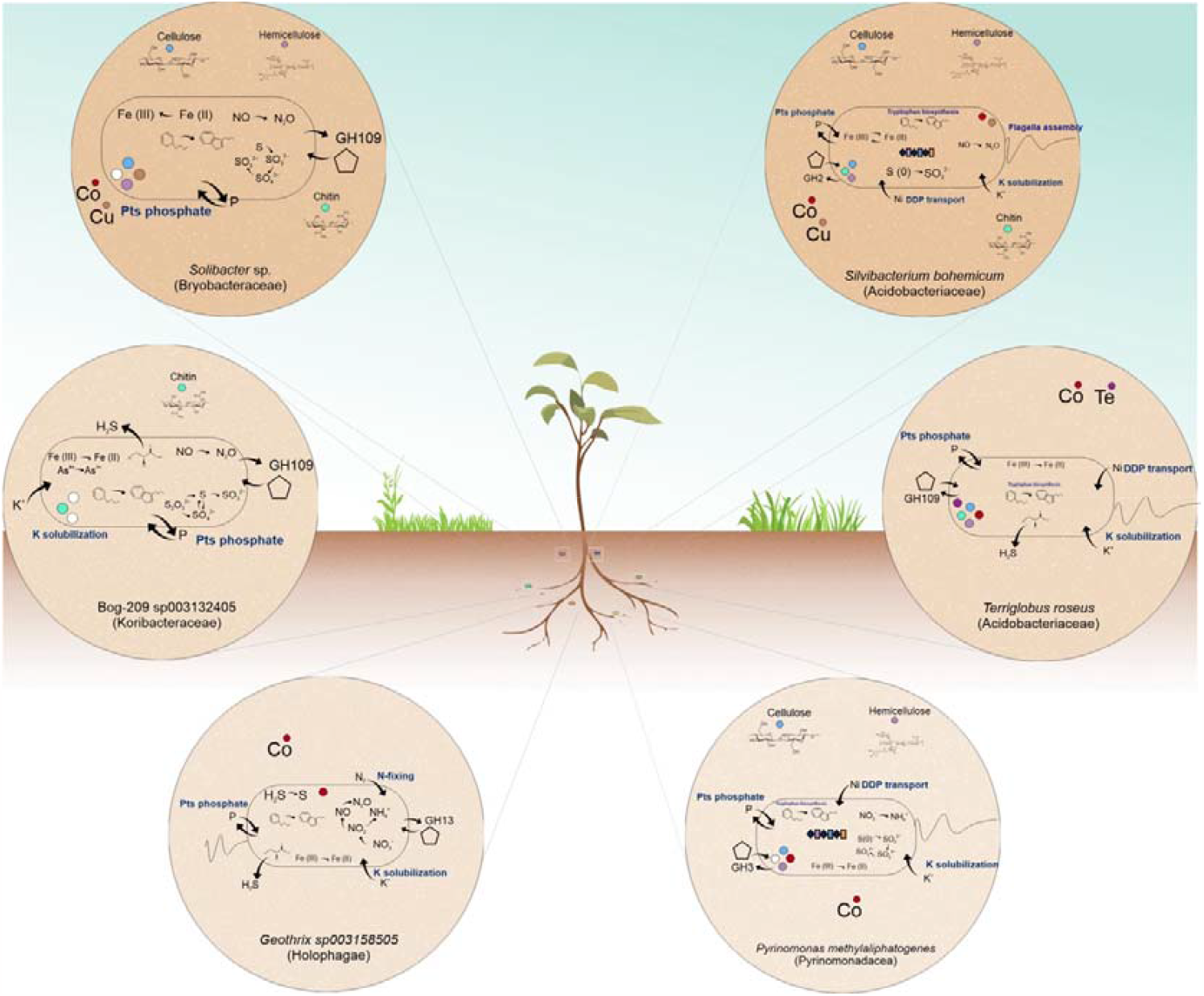
Schematic illustration highlighting some representatives of Acidobacteriota families involved in plant-microbe interactions and their potential influence on the biogeochemical processes contributing to soil functioning. Distinct bacterial groups are distinguished by various circle augmentations, while biogeochemical reactions are depicted within each cell. The ability to break down complex carbon compounds and withstand heavy metals is visually represented through circle colors. Curved lines illustrate transformations occurring in the periplasm or involving membrane-bound processes, while outer pentagon shapes represent extracellular CAZYmes. The lower part of the figure showed general microbiome services and plant-specific traits.

We also observed that the solubilization of insoluble potassium (K) is a common trait among these representatives, with the ability to convert it into a soluble form suitable for plant growth (Etesami et al., 2017). Additionally, we found that some members of Acidobacteriota are capable of surviving in environments contaminated with heavy metals, suggesting their potential as bioremediation agents. Furthermore, the presence of CRISPR loci and other antiviral mechanisms in *Acidobacteriaceae* and *Pyrinomonadaceae* members may provide a selective advantage in the interaction between bacteria and phages in the soil environment (Doron et al., 2018). In the biogeochemical context, we observed that many members of Acidobacteriota are involved in nitrogen transformation processes such as ammonification, nitrification, and biological nitrogen fixation, as well as sulfur-related processes including sulfate reduction, sulfur oxidation, and sulfide oxidation (Fig. 6). Although it is challenging to establish a direct effect of these bacterial plant growth-promoting traits *in vitro* (Yoneda et al., 2021), either due to the lack of understanding of their mechanisms or the difficulty of isolating and cultivating in laboratory conditions, the transformations performed by Acidobacteriota in oxidizing essential elements for plant growth may improve overall plant growth. Altogether, this study highlights the importance of this intriguing phylum and opens new avenues for future research aimed at exploring specific Acidobacteriota taxa and their implications in plant interactions for applications in agriculture and environmental sustainability.

## Acknowledgements

The authors expressed their gratitude to the technical support team of the Cluster at Universidade Federal de Viçosa. We expressed our appreciation to Felipe Xavier for providing valuable artistic insights into Figure 6 of this paper.

## Contributions

Conceptualization: OG, and MF. Data curation: OG, AF, ST, TF, LA. Funding acquisition: MF. Investigation: OG, AF, ST, TF, LA, CC. Methodology: OG. Project administration: MF, Resources: OG and MF. Supervision: MF and CC. Visualization: OG. Writing – original draft: OG, AF, ST, TF, LA. Writing – review & editing: MF and CC.

