## Supplementary note for "Insights into Plant Interactions and the Biogeochemical Role of the Globally Widespread Acidobacteriota Phylum"

**Supplementary Note 1**

Metagenomic data has revealed intriguing findings regarding the diversity of Acidobacteriota in various environments [1–3]. Notably, dissimilarity in sulfate reduction pathways (dsrABC) was not detected in the soil Acidobacteriota, specifically within subdivision 1 and 3 strains, contradicting the initial genome data reports [4]. Conversely, further analysis employing metagenomic data shed light on the presence of genes encoding DsrD within Acidobacteria members (Anantharaman et al. 2018) Likewise, in marine sediment environments, Acidobacteriota species like *Candidatus* *Sulfomarinibacter* (class Thermoanaerobaculia, subdivision 23) and *Ca*. *Polarisedimenticola* (subdivision 22) have been identified to harbor versatile respiratory pathways [3]. These pathways encompass a range of possibilities, including oxygen, nitrous oxide, metal-oxide, tetrathionate, sulfur, and sulfite/sulfate respiration, and potentially even sulfur disproportionation [3]. Remarkably, our dataset also corroborated the presence of these classes, with a pronounced dissimilarity in sulfate reduction predominately linked to aquatic environments.

As previously reported, we did not find any evidence of nitrification capable Acidobacteriota strains [4–6]. In addition, our study has unearthed a cohort of Acidobacteriota members playing a key role in nitrogen fixation. Within this context, we have identified and characterized 17 MAGs exclusively affiliated with the *Acidobacteriaceae* and *Holophagae* families. Interestingly, most of these genera have not been taxonomically classified yet. We did not observe any direct correlation, beyond the family level, between these genera and their involvement in nitrogen fixation. Therefore, we hypothesized that this phenomenon could potentially be attributed to horizontal gene transfer events. However, due to the inherent limitations of MAGs, we were unable to definitively determine the origin of these nitrogen-fixing capabilities. Further investigations are needed to fully understand the genetic basis and evolutionary dynamics of nitrogen fixation in these Acidobacteriota genera. Furthermore, our findings build upon prior evidence indicating nitrogen fixation capabilities within the Acidobacteriota [7].

We have identified a set of MAGs within the Acidobacteriota, all encoding a membrane-bound respiratory nitrate reductase. Among these MAGs, the most prevalent orders were Acidobacteriae, Thermoanaerobaculia, and Blastocatellia. This functionality is not restricted to a single habitat, encompassing prominent members from both soil and aquatic environments, such as *Granulicella*, *Acidobacterium*, *Koribacter*, and even the sulfate-reducing bacterium *Sulfotelmatobacter* [3]. This role is particularly associated with the *Pyrinomonadaceae* family within Blastocatellia [8], and the *Thermoanaerobaculaceae* family within Thermoanaerobaculia. This family have only one formally described member so far, characterized as a Gram-negative, thermophilic, and neutrophilic anaerobic bacterium [9]. Intriguingly, our study has revealed the presence of *napAB*|*narGH* encoding genes in the genome of *Thermoanaerobaculum* *aquaticum*, despite the absence of observed utilization of nitrate and nitrite as electron acceptors, as reported for strain MP-01T [10]

Notably, consistent with the overall dataset, our analysis did not uncover any Acidobacteriae members encoding *nosDZ* genes associated with Nitrous oxide reduction. This function appears to be primarily linked to the *Pyrinomonadaceae* family within Blastocatellia, Thermoanaerobaculia, and Luteitalea, a member of subdivision 6 Acidobacteriota isolated from temperate grassland soil [11]. To summarize, our findings provide compelling evidence of Acidobacteria's active participation in pivotal nitrogen cycle processes, including nitrogen fixation and denitrification.
