## Supplementary Figures for "Insights into Plant Interactions and the Biogeochemical Role of the Globally Widespread Acidobacteriota Phylum"

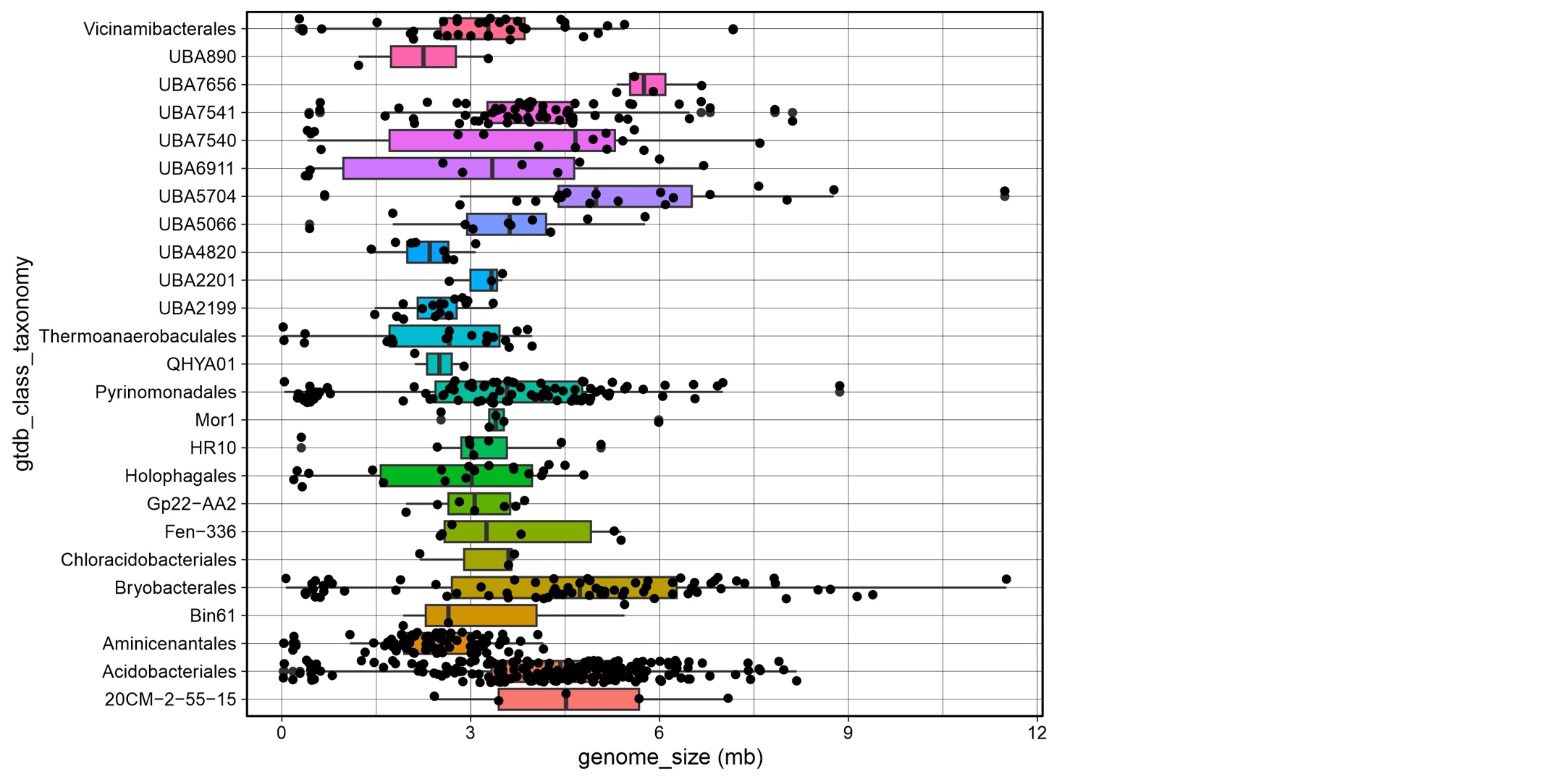

**Fig S1. Boxplot depicting the genome size of Acidobacteriota families mapped in the dataset**. The y-axis displays the taxonomic family of Acidobacteriota, while the y-axis represents genome size in megabases (Mb).

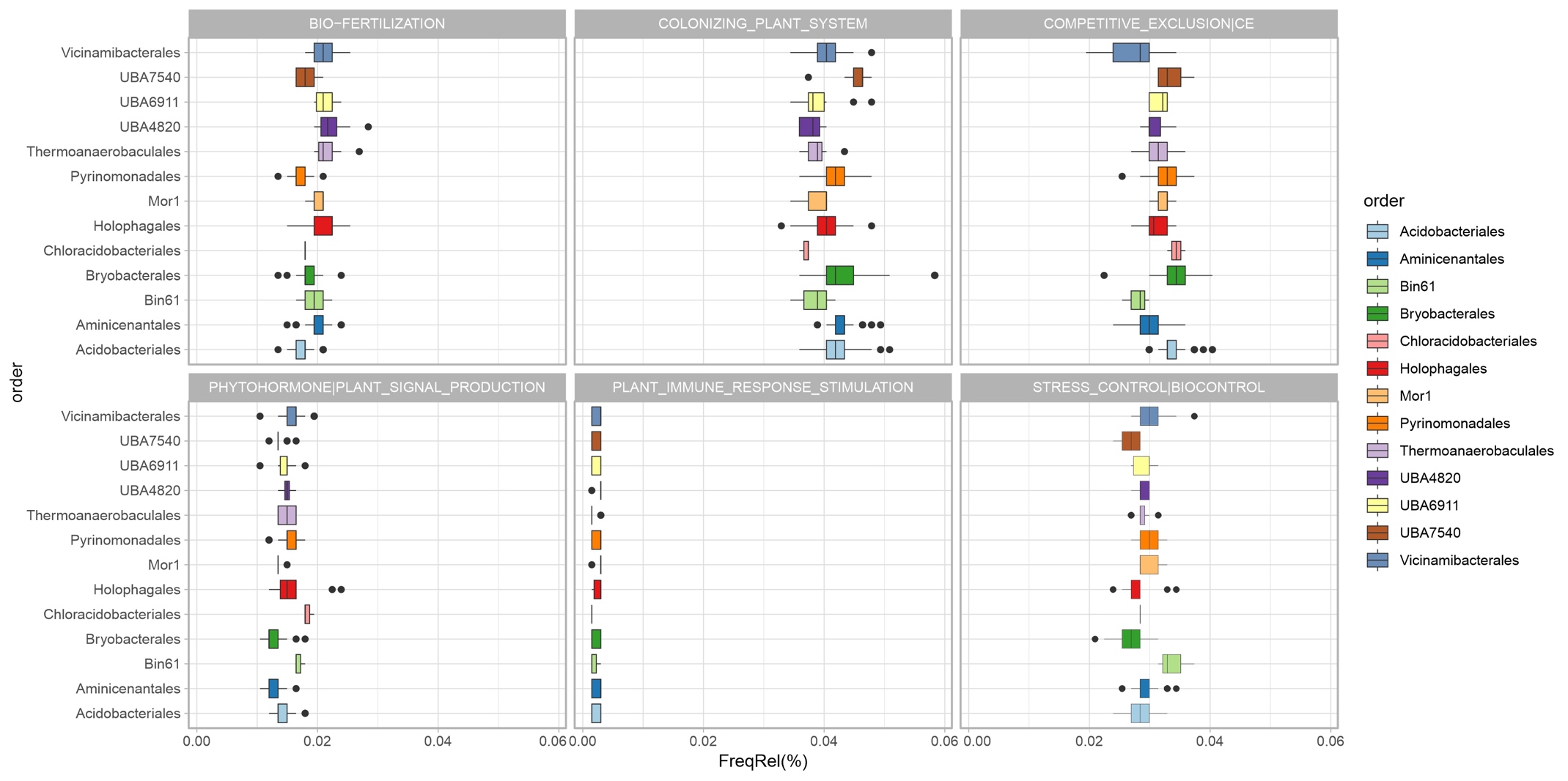

**Fig S2.	Relative frequency of plant growth-promoting gene classes identified within the Acidobacteriota order**

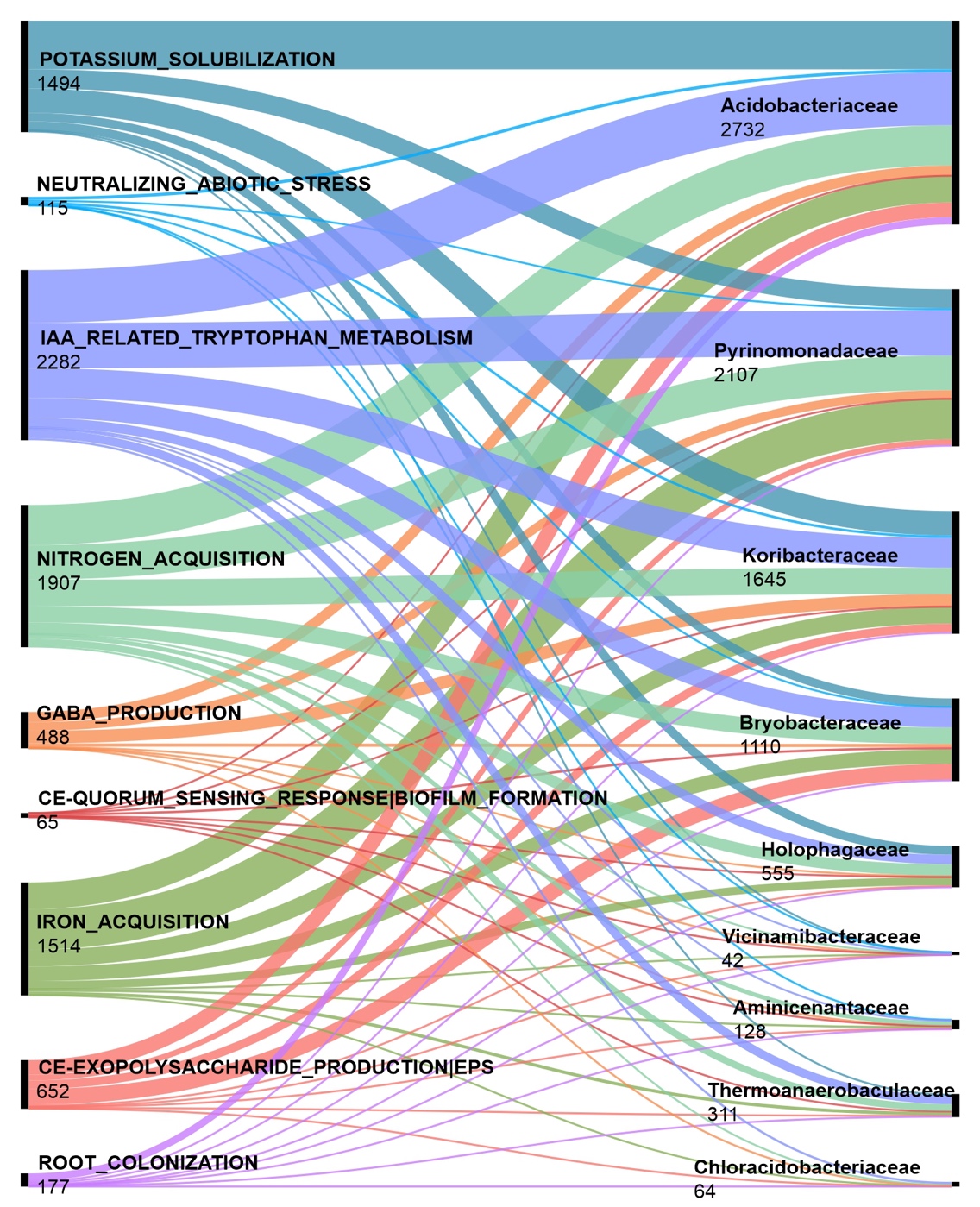

**Fig S3. The distribution of 86 plant growth-promoting gene classes across Acidobacteriota families**. The numbers below each class and family indicate the number of hits found.

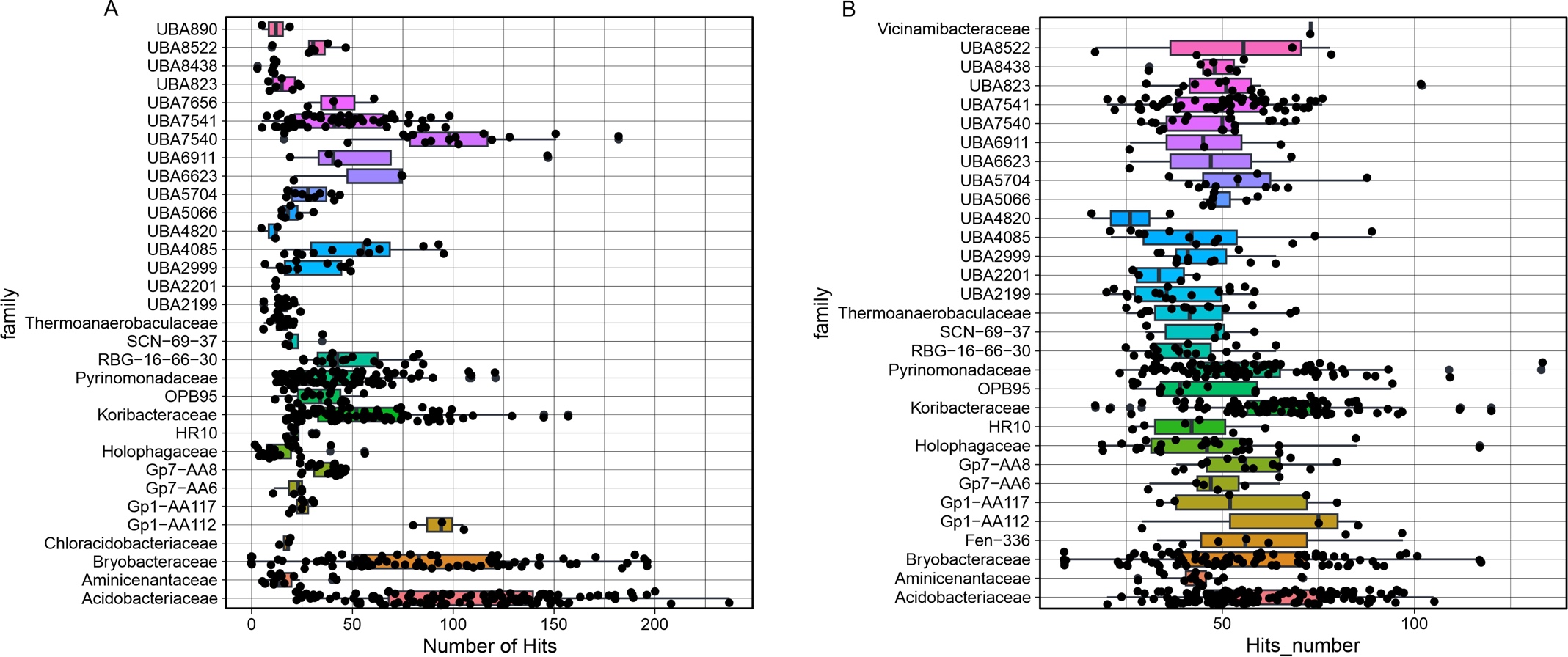

**Fig. S4. Potential of Acidobacteriota to act in plant polymer degradation.** (A) Box plot showing the number of Carbohydrate Active Enzymes (CAZymes) across Acidobacteriota family. (B) A) Box plot showing the number of peptidases across Acidobacteriota family.

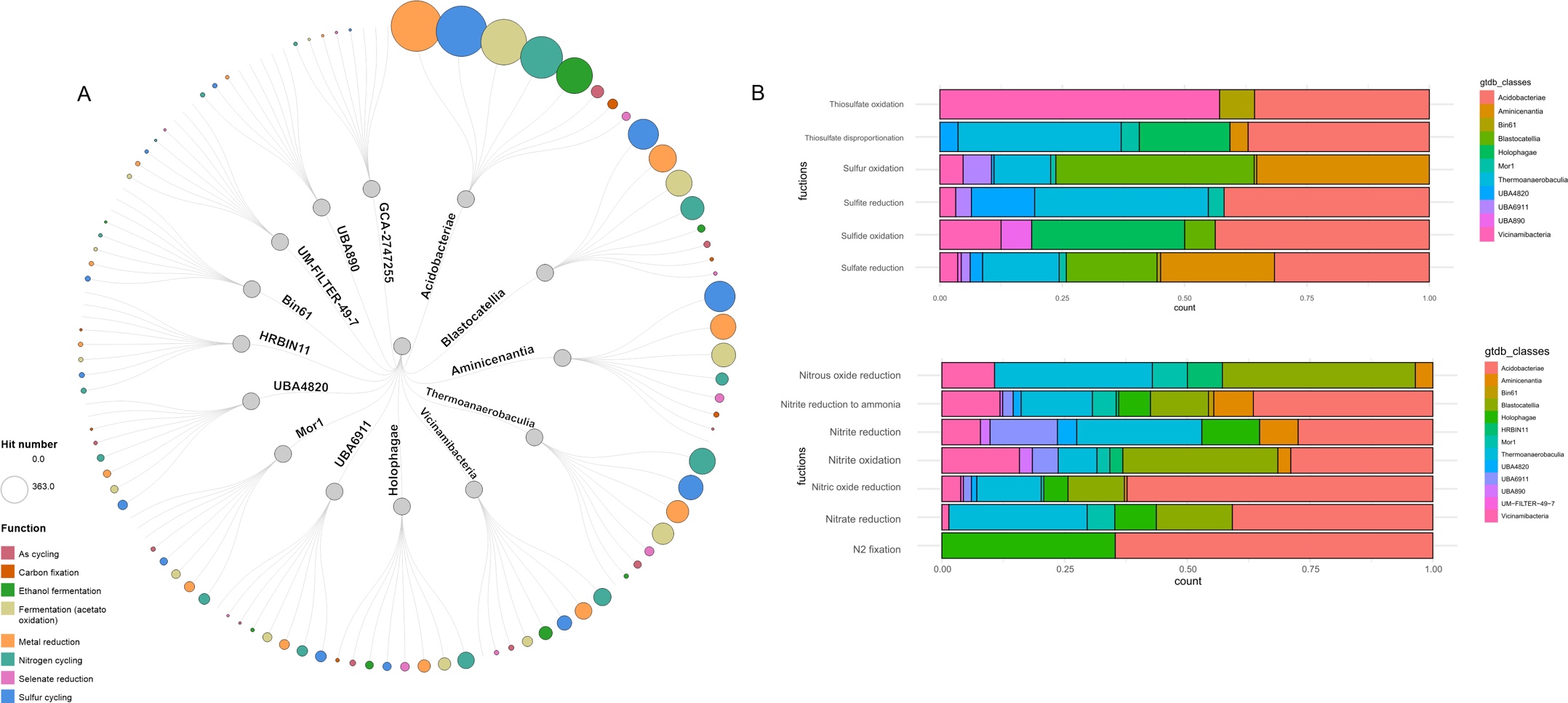

**Fig. S5. The distribution of metabolic cycles among 14 representative Acidobacteriota families**. (A) A circular dendrogram is presented, depicting nine biogeochemical cycles across the families. Each cycle is color-coded, and the size of the circles represents the number of hits. (B) The composition of functions related to the sulfur cycle (top-graph) and nitrogen cycle (bellow-graph) within each family is displayed.

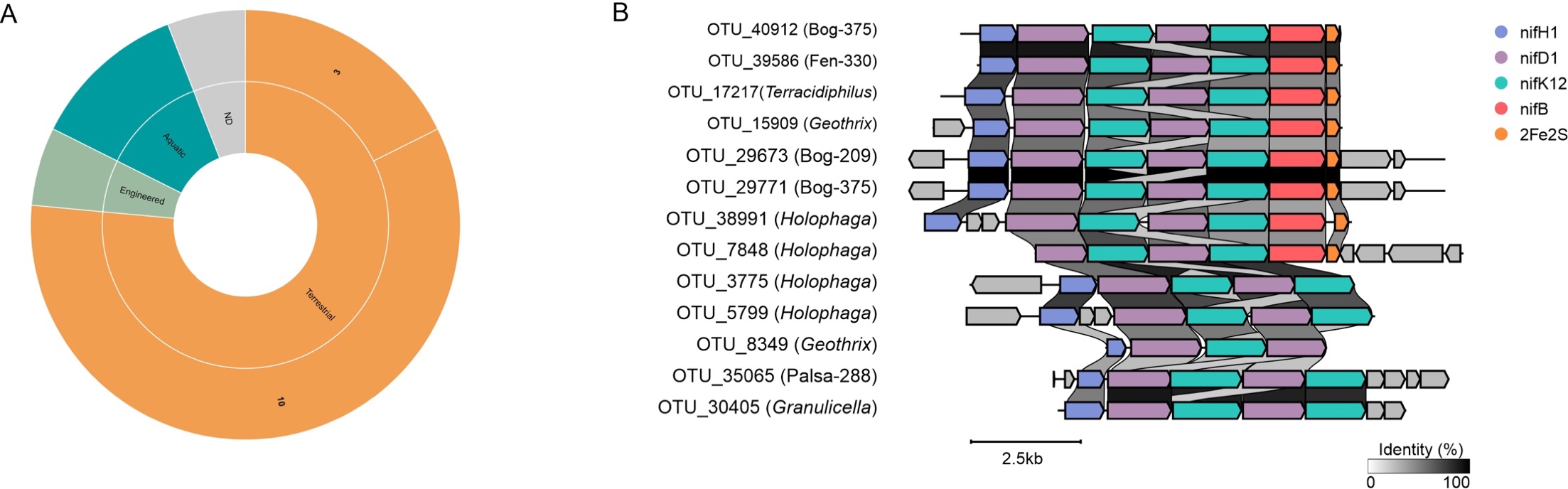

**Fig. S6. The role of 17 metagenome-assembled genomes (MAGs) belonging to Acidobacteriota in nitrogen fixation.** (A) The environmental information of the MAGs is presented, indicating their distribution in terrestrial environments (15), aquatic environments (2), and engineered environments. (B) A syntenic analysis is performed on 13 *nif*-clusters mapped across the MAGs. Genes within the clusters are color-coded based on their function, providing insights into their functional similarities.
